# Costs of resistance limit the effectiveness of cooperation enforcement

**DOI:** 10.1101/2023.08.29.555259

**Authors:** M.E. Miller, S. Sidell, E.A. Ostrowski

**Affiliations:** School of Natural Sciences, Massey University, Auckland, New Zealand; Department of Biology and Biochemistry, University of Houston, Houston, TX, USA

**Keywords:** Cooperation, cheating, experimental evolution, trade-offs, coevolutionary arms race, social conflict

## Abstract

Cooperative groups are susceptible to invasion by cheaters that reap the benefits but fail to pay the costs. Both theory and experimental work have shown that cheating can select for counter-adaptations to resist cheating. But then why is cheating so common? One key hypothesis is that trade-offs prevent resistors from taking over, but evidence to support the trade-off model is lacking. Here we evolved resistance to different cheaters and tested for trade-offs. Improvements against one cheater frequently entailed correlated improvements against novel cheaters. However, direct responses to selection were typically stronger than correlated responses to selection, resulting in a pattern of local adaptation of resistance. Control populations, evolved in the absence of a cheater, showed improvements in spore germination, whereas cheater-evolved populations did not, suggesting that the evolution of resistance retards or prevents other fitness improvements. Taken together, our findings suggest that, although cheater resistance can evolve rapidly, it may also involve subtle trade-offs that can help to explain the maintenance of polymorphism in cheating and resistance in nature.

## Introduction

Cooperative societies can be exploited by individuals that benefit from cooperation without paying their fair share of its cost, termed cheating (Ghoul et al 2014, Leeks et al 2021; Jones et al. 2015; West et al. 2007). The fitness benefits of cheating can allow cheaters to invade and, in some cases, threaten the existence of cooperative societies (Fiegna and Velicer 2003; Rainey and Rainey 2003; Leeks 2021; MacLean Heredity 2007; Hardin 1968). Thus, a key question is how cooperative societies prevent the invasion and spread of cheaters.

One way to limit the spread of cheating is to cooperate only with kin, which ensures that the beneficiaries of cooperation are likely to pass on the altruist’s genes (Hamilton 1964). However, not all societies are composed of relatives, and kin selection alone may not be sufficiently strong to suppress cheating (Frank 1995; Van Dyken et al. 2011). An alternative hypothesis is that selfish behaviors might be countered by evolutionary changes to suppress or resist their effects— termed enforcement (Frank 1995, 2003; El Mouden et al. 2010; Ågren et al. 2019). Enforcement of cooperation is common in eusocial insects, where workers can behave selfishly by laying their own eggs, but other workers can punish these actions by attacking and killing the individual or the eggs (Ratnieks and Wenseleers 2005). In microbes, resistance evolves rapidly to suppress selfish mutants, suggesting that the counter-evolution of cheating resistance is a plausible mechanism to limit the long-term success of cheaters and stabilize cooperation (Khare et al. 2009; Kümmerli et al. 2010; Manhes and Velicer 2011; Hollis 2012; Levin et al. 2015).

While suppression seems plausible, the success of resistance in stemming cheating is not clear. Several questions remain. How readily does resistance evolve? Can different types of selfishness be suppressed, or are some ways easier to counter than others? Are there costs of being resistant, and how do those costs influence the success of enforcement? Drawing on the concepts from host-pathogen co-evolution (Frank 1992; Brown 2003; Burdon and Thrall 2003; Hall et al. 2011; Koskella et al. 2012; Brown and Rant 2013; Scanlan et al. 2015), we can identify two distinct types of costs of resistance, which need not be mutually exclusive: (1) host-specialisation trade-offs (HSTOs), where improvements against one cheater are detrimental for the ability to suppress a different cheater, (2) life-history trade-offs (LHTOs), where resistance improves one fitness component but comes at the cost to others, including adaptations to the abiotic or asocial environment (Barrett et al. 2011; Scanlan et al. 2015). Both HSTOs and LHTOs have important consequences for the evolutionary dynamics of cooperation. HSTOs could lead to an ongoing co-evolutionary arms race, where resistors are never fully adapted to any novel cheater they encounter, and populations can evolve divergently in response to the unique cheaters they experience. HSTOs can thus result in local adaptation and a matching of cheating and resistance alleles within populations (Kaltz and Shykoff 1998; Nuismer 2006; Kraemer and Kassen 2015; Butaitė et al. 2021) LHTOs might lead to balanced polymorphisms or trench warfare dynamics, where cheating and resistance alleles are both maintained long-term in populations (Barrett et al. 2011; Ostrowski et al. 2015).

Here we address the existence of cheater-resistance trade-offs using experimental evolution and a model system for cooperation and conflict. The social amoeba *Dictyostelium discoideum* is a haploid, eukaryotic amoeba that lives in the forest soil of North America and east Asia. When cell densities are high and starvation is imminent, the amoebae aggregate and take on a multicellular form with differentiated cell types. Starving cells aggregate, form a mound, and then transform into a multicellular, migratory slug with differentiated cell types. Following a period of migration, the slug transforms into a multicellular fruiting body. The cells in the anterior form a rigid stalk and then vacuolize and die. The remaining cells rise to the top of the stalk and form environmentally resistant spores, which can disperse and germinate to form single-celled amoebae again. Stalk formation is altruistic, in that stalk cells give up their lives, ostensibly because their death provides a fitness benefit to the spores. Spore-stalk differentiation is akin to germline-soma differentiation that occurs in other forms of complex multicellularity.

Conflict can arise at the transition to multicellularity because genetically different strains can co-aggregate, resulting in a chimeric organism, and stalk and spore cells have drastically different fitness outcomes vs.(survival vs certain death; Ostrowski 2019). This combination of ample genetic variation and strong selective consequences of the different cell fates allows selection to favor genotypes that adopt the spore cell fate and avoid the stalk cell fate. Indeed, studies of naturally occurring variants and laboratory mutants have demonstrated that such spore-preferring strains exist and arise readily (Strassmann et al. 2000; Santorelli et al. 2008; Buttery et al. 2009). Since then, substantial work has been aimed at understanding whether and how the organism maintains altruistic stalk formation when its life cycle seems prone to promoting selfishness. The answers to these questions have broad implications for how complex life forms evolve and whether that complexity can be maintained in the face of selection that favors individual success over that of the group.

We used laboratory experimental evolution to select for resistance in response to four genotypically distinct cheaters (Fig. 1A). Each cheater forms a disproportionate share (>50%) of the spores when co-developed at equal starting ratios (50:50) with its wild-type progenitor AX4 (Fig. 2). Following multiple rounds of growth and co-development in the presence of a cheater, we tested the evolved populations for their ability to suppress cheating by their focal cheater, termed direct resistance (Fig. 1C). To test for HSTOs, we asked whether evolved populations can suppress unfamiliar cheaters, those to which they were not exposed, termed cross-resistance (Fig. 1C). We hypothesised that cross-resistance would be limited, leading to strains that are locally adapted to the strains they encountered during their evolution. We furthermore hypothesised that improvements during the social stage (e.g., against a particular competitor) might come at a cost to fitness at other stages of the life cycle (LHTOs), and therefore we asked how well resistant populations carry out other functions necessary for fitness in this environment. Together, these processes might limit the extent and duration of successful resistance.

**Figure 1.**
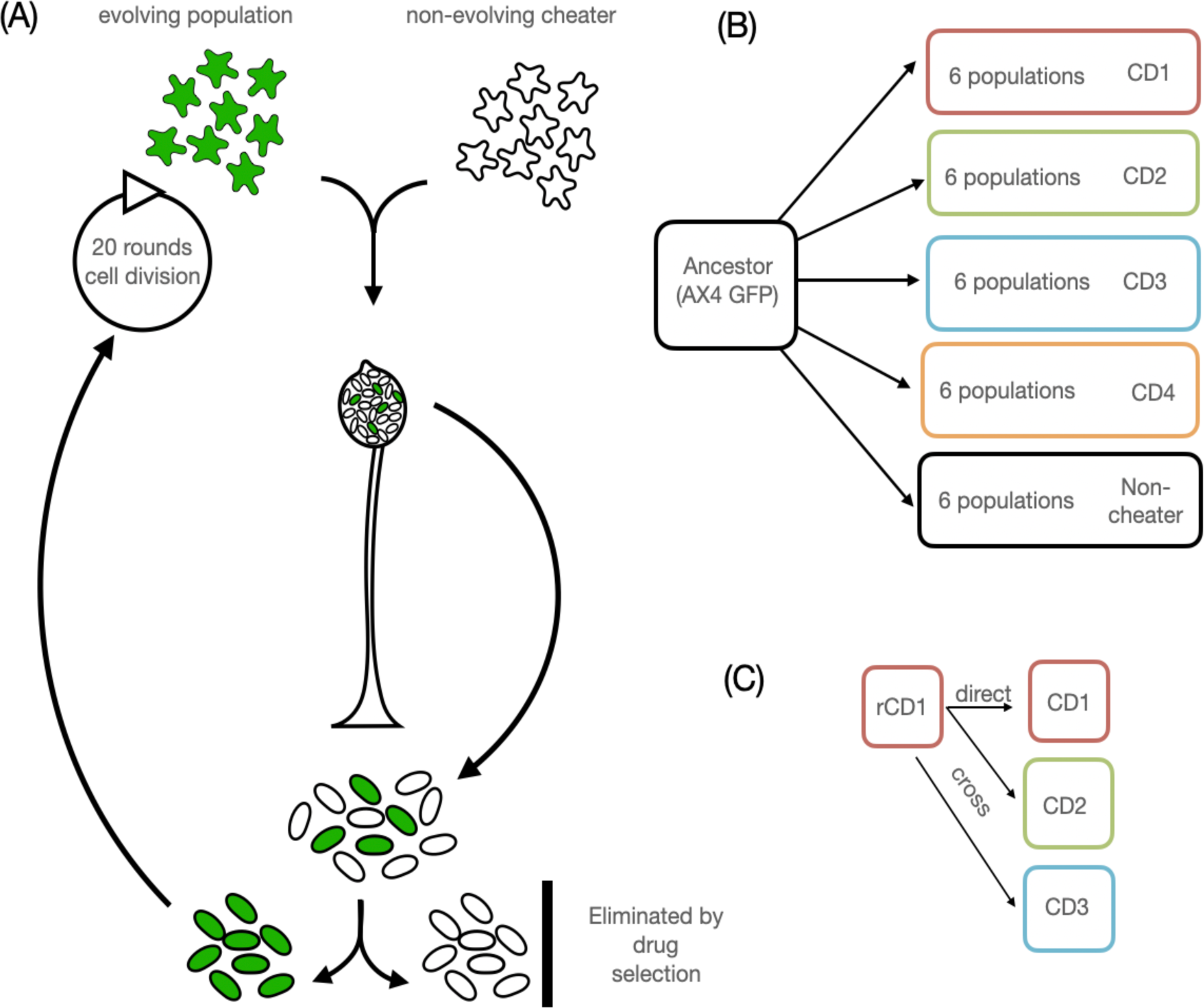
Selection for cheater resistant strains. (A) Selection for cheating resistance, where cells of the evolving population (GFP-labeled) are co-developed with cells of the cheater strain. Each round, the cheater is eliminated by drug selection. The surviving cells are regrown, combined again with cells of the cheater, and re-developed. (B) Six replicate populations were evolved with each of the CD1-CD4 strains, as well as a non-cheating control (AX4). All populations underwent 15 rounds of social development, with approximately 20 generations per round. (C) All-against-all design. All evolved strains (e.g., ‘rCD1.1’) were tested against the cheater they faced during their evolution (‘direct resistance’), the other three cheaters (‘cross resistance’), and non-cheating parental strain.

**Figure 2.**
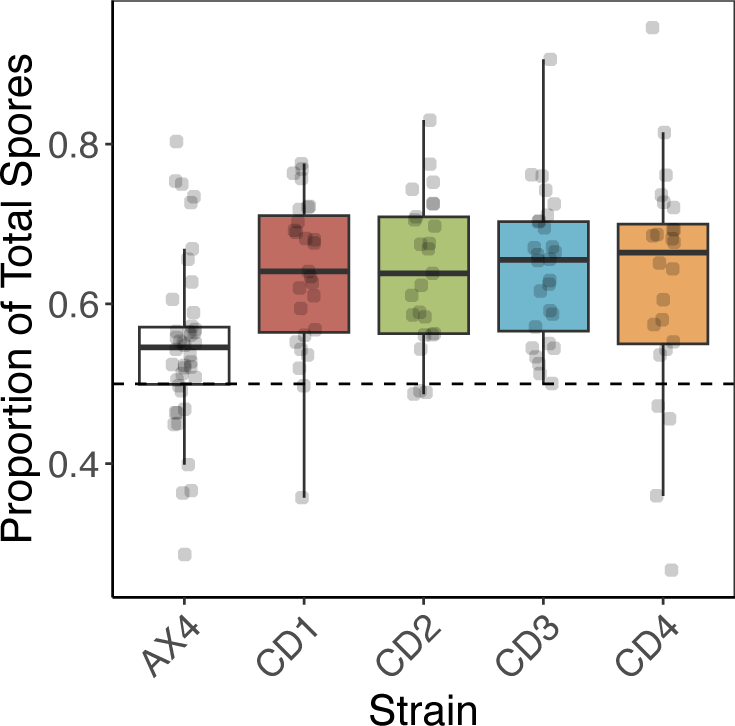
CD strains form more than their fair share of spores when co-developed with their parent strain, AX4. Each dot represents the results of single mix experiment between the indicated strain and AX4-GFP. The dotted line indicates the expected frequency of the spores if development is perfectly fair. AX4 is the control mix, consisting of the parental strain AX4 co-developed with AX4-GFP.

## Methods

### Resistance Evolution Experiment

At the start of the experiment, we combined cells of the ancestor (AX4-GFP) in equal proportions with cells of one of the four cheaters (CD1-CD4) or the non-cheating progenitor (AX4) and allowed them to fruit together. For each cheater, we initiated six replicate populations that were combined with one of the cheaters, resulting in 30 evolving populations (= 5 social partners x 6 replicates per partner; Fig. 1B). After each round of fruiting, we collected the spores, preserved an aliquot in the freezer, and then germinated the remaining spores at a density of 5×10^4^ spores/ml in HL5 medium, which was supplemented with 30 μg/ml of G418 to remove either the drug-sensitive cheater or the non-cheating wildtype strain (in the case of the non-cheater control treatments). Once the surviving cells had regrown, we repeated the process, combining the survivors of the previous round again with their focal cheater and allowing them to fruit together. We grew the cheater strain from frozen stocks at the start of each round, and thus the populations were selected for improvements against an evolutionarily static target genotype.

We carried out 15 rounds of growth and fruiting in the presence of their focal cheater strain. Each round consisted of a growth phase, where the surviving cells from the previous round were re-grown to high density for approximately one week, equal to roughly 20 generations, followed by a round of development with their target genotype. In total, the experiment encompassed approximately 300 generations of evolution (∼3 doublings/day x 7 days/round x 15 rounds). At the end of the experiment, we tested the evolved populations for their resistance to the social competitor they experienced during their evolution, their cross-resistance to unfamiliar competitors, and their evolutionary change in other fitness-related traits, including their doubling time in the liquid medium, spore germination, and total spore production. We use these results to assess the extent to which resistance evolves, whether it confers cross-resistance (HSTOs), and costs of resistance (LHTOs).

#### Strains

The four cheater strains, CD1 to CD4, were provided by Adam Kuspa and Chris Dinh, Baylor College of Medicine. These strains were generated by restriction enzyme-mediated insertional (REMI) mutagenesis of the standard lab strain, AX4. These particular strains were isolated from a selection experiment where mutant pools of 600-800 strains were propagated through repeated rounds of growth and chimeric co-development, a process that can enrich for strains that form more spores than their competitors. The plasmid insertion site in each strain was subsequently identified through plasmid rescue or whole genome re-sequencing (Table S1).

#### Culture conditions

Frozen spore stocks were spotted on SM agar plates (Formedium Ltd, 2% agar) with 400 μl *Klebsiella pneumoniae* as a food source. Once fruiting bodies had formed, spores were inoculated into Petri dishes containing 10 ml of HL5 medium (Formedium Ltd), supplemented with PSV (10 μg/ml penicillin, 50 μg/ml streptomycin sulfate, 60 ng/ml cyanocobalamin, and 20 ng/ml folate). GFP-labeled strains received 10 μg/ml G418. Once cells reached confluence, the cultures were transferred to flasks and maintained with shaking at 180 RPM at 22 °C. Cultures were maintained in exponential phase through daily dilutions into fresh media for approximately one week prior to each round of development.

#### Chimeric Multicellular Development

For chimeric co-development, cells were harvested during mid-exponential growth, washed twice in cold KK_2_ buffer (per L: KH_2_PO_4_ 2.25 g, K_2_HPO_4_ 0.67 g), and resuspended at 1×10^8^ cells/ml in KK_2_. GFP-labeled cells were combined with their non-labeled competitor at a 1:1 ratio and aliquots corresponding to 2×10^7^ cells were deposited in a 6×6 square on 47 mm 0.45 μm nitrocellulose filters, resulting in a density of ∼5.8×10^5^ cells/cm. The filters were placed on top of Pall filter pads with 1.5 ml PDF (20.1 mM KCl, 5.3 mM MgCl_2_·6H_2_O, 9.2 mM K_2_HPO_4_, 13.2 mM KH_2_PO_4_, 0.5 g/L streptomycin sulfate) in 6-cm Petri dishes. As a control, all strains were simultaneously developed clonally (that is, without a partner strain). Plates were incubated at 22°C in humid chambers with overhead light. To retrieve spores from the fruiting bodies, the filters were transferred after 48 h to 50 ml Falcon tubes containing 5 ml detergent (KK_2_ buffer, 0.1% IGEPAL, 20 mM EDTA), vortexed, and counted using a hemacytometer or cell counter (Countess II, ThermoFisher). The percentage of cells expressing GFP was determined using flow cytometry (Accuri C6, BD).

#### Germination Assays

Spores were spotted from frozen stocks onto SM agar plates spread with 400 μl of *K. pneumoniae*. Once fruiting bodies formed, spores were collected and diluted to 5×10^4^/ml in HL5 and placed in wells of a 96-well plate. The wells were photographed immediately after plating and then again after incubation for 12 hours. Germination efficiency was calculated as one minus the proportion of spores present after 12 hours incubation.

#### Doubling Time

To estimate the doubling time, changes in cell density of shaking cultures were estimated using cell counts during exponential growth. Doubling time (in hours) was estimated as the number of hours between the two cell counts, divided by the number of doublings that took place in that time interval.

#### Sporulation efficiency

The sporulation efficiency of each strain, defined as the fraction of starting cells that form spores, was estimated using the clonal development filters from the co-development assays. After 48 hours, the filters were transferred to 50 ml Falcon tubes containing 5 ml detergent to lyse any remaining cells, and an aliquot was counted using a hemocytometer or an automated cell counter (Countess II, ThermoFisher).

#### Statistical analyses

We used generalized linear mixed effect models to test for resistance. Statistical analyses were conducted in R (R Core Team 2018) with the packages glmmTMB for model building and emmeans for posthoc testing (Brooks et al. 2017; Lenth 2023). Each strain’s proportion of the spores in chimeric development with another strain was modelled as a function of the strain it was being tested against (=current social partner), its evolutionary history (=historical social partner), the interaction between the two, the mix type (‘direct-’ or ‘cross-’ resistance; see Fig. 1C), and the experiment date (=block). We used a beta-family distribution and logit link function. For random effects, significance was determined by comparing the fits of models with and without factors of interest, and non-significant factors were dropped. We removed any data points if either of the two strains being tested failed to develop in the clonal controls.

#### Whole-Genome Resequencing and Variant Calling

Detailed methods for whole genome sequencing are provided in the supplement. Briefly, genomic DNA was purified from nuclei using phenol-chloroform extraction. Libraries were prepared using the NexteraXT kit, with 0.25x volumes prior to size selection using SPRIselect beads (Becker Coulter B23319) for a target size of 300-500 bp. Sequencing was carried by Admera Health. Raw sequencing reads were mapped to the chromosomal sequences of the AX4 reference genome using BWA-MEM, and SNP calling was performed using GATK with a hard filter; filtering parameters are described in the supplement. Since *D. discoideum* is haploid, we required that at least 80% of the reads covering a site showed the alternate allele. We also removed variant sites if no strain showed strong evidence for the reference allele (defined as a minimum of 80% reads covering the site showing the reference nucleotide), as these sites potentially represent poorly mapped regions or errors in the reference genome.

## Results

### Mutants obtain more than their fair share of spores

We first tested whether the four strains (CD1-CD4) isolated from the initial selection for cheating do in fact form a disproportionate share of the spores when co-developed with their parental strain, AX4, at starting ratio of 1:1. These results show that the CD strains do form more than their fair share of spores (all *P*<0.001; Fig. 2).

### Evolved Strains Show Improvements in Spore Equity

After 15 rounds of co-development, the evolved populations were tested for their ability to obtain spores when co-developed with their evolution partner strain. Most populations showed a higher mean representation in the spores than their ancestor (Fig. 3). In all four CD treatments, the evolved populations show significant improvements against their focal cheater compared to their ancestor (emmeans contrasts based on generalized linear model; all *P*<0.05; Table 1). However, the control populations that evolved without a cheater (i.e., the AX4-evolved population) did not show significant improvement relative to the ancestor (Table 1).

**Figure 3.**
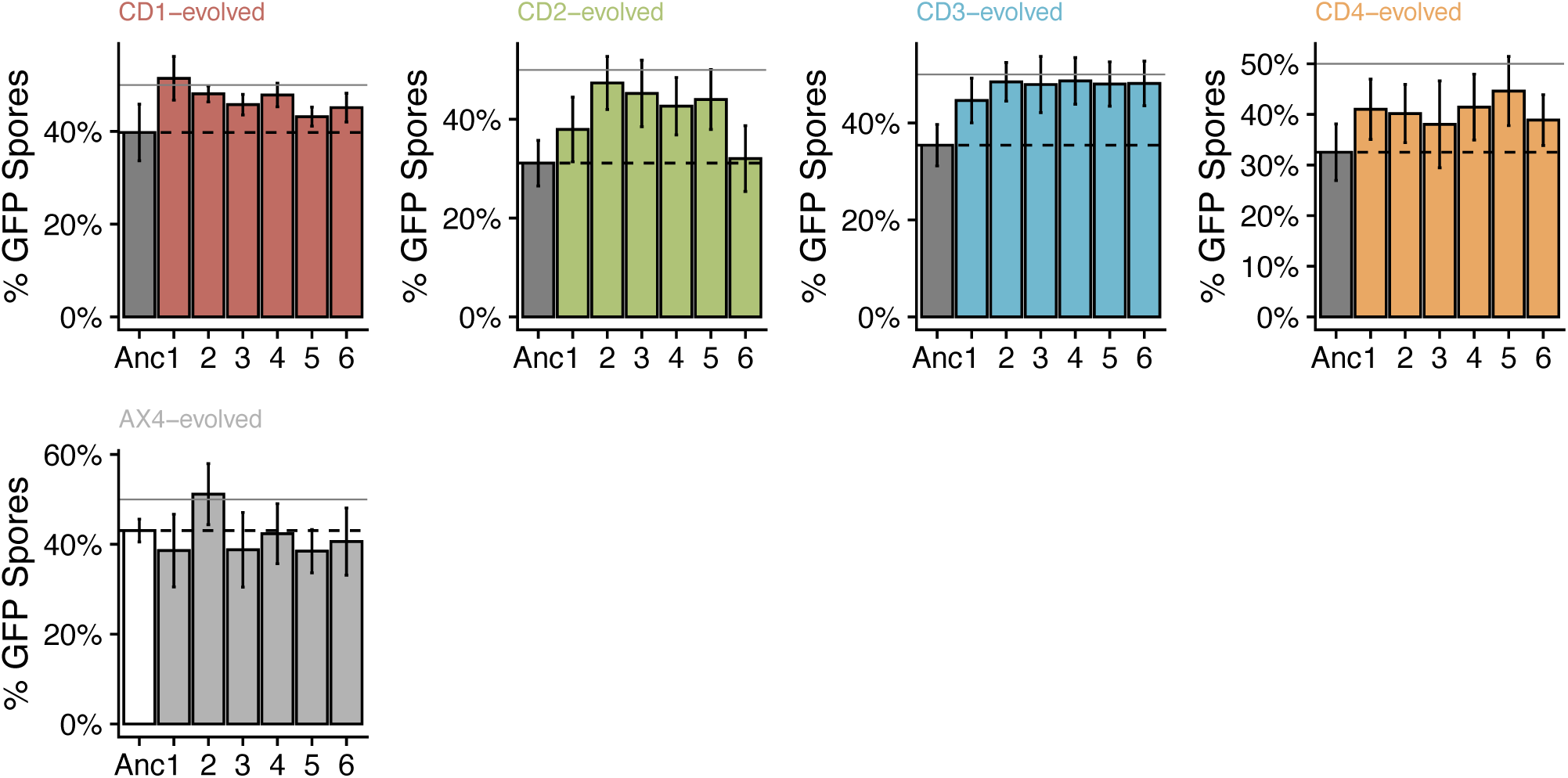
Evolved populations show quantitative improvements against their respective cheaters. Each box shows the mean percentage of spores obtained by the indicated population (ancestor or evolved populations 1-6) following chimeric development with the cheater they evolved with. Dotted reference lines denote the performance of the ancestor against that cheater, which is always below 50%. Solid lines indicate the expected (=50%) percentage of spores if fully resistant to cheating. Most populations lie above the dotted line, indicating some evolutionary improvement, yet below the solid line, indicating that they are not fully resistant.

**Table 1.** Contrasts comparing performance of evolved populations to their ancestor for each cheater (CD1-4) or the non-cheating control (AX4).

| Contrast | Estimate | t | df | P |
| --- | --- | --- | --- | --- |
| Anc vs CD1-evolved | -0.31 | -2.17 | 36 | 0.0368 |
| Anc vs CD2-evolved | -0.51 | -2.03 | 36 | 0.0497 |
| Anc vs CD3-evolved | -0.52 | -4.23 | 36 | 0.0002 |
| Anc vs CD4-evolved* | -0.407 | -3.43 | 25 | 0.0021 |
| Anc vs AX4-evolved | -0.079 | -0.54 | 35 | 0.5921 |

\*Experiment date was modelled as a fixed effect to achieve model convergence.

We also isolated single clones from each population and tested each one’s ability to resist cheating relative to its ancestor (Fig. 4). Because the evolved populations can be genetically diverse, each isolated clone will not necessarily show the same level of resistance as the population as a whole. Overall, the majority are more resistant than their ancestor to their focal cheater. For CD1-, CD3-, and AX4-evolved populations, the clones were significantly better than their ancestor at obtaining spores when paired with their respective partners (emmeans contrasts; all *P*<0.05). CD2-evolved populations were not significantly more resistant than their ancestor, but this result was driven by one clone (from population rCD2.6) that performed especially poorly; excluding this one clone showed that the remaining 5 populations were significantly more resistant than their ancestor. Finally, clones isolated from the CD4-evolved populations were not significantly resistant to CD4 (Table 2, *P*=0.67), with 4 of 6 clones showing a lower mean performance than the ancestor against CD4. Unlike CD2-evolved strains, this pattern could not be explained by just one strain. Moreover, none of the CD4-evolved isolates were significantly better than their ancestor (all *P*>0.05). We exclude these strains in the analyses below, since the question of whether resistance trade-offs with other traits is predicated on it having evolved in the first place. We discuss the apparent lack of resistance in these isolates in more detail in the supplement.

**Figure 4.**
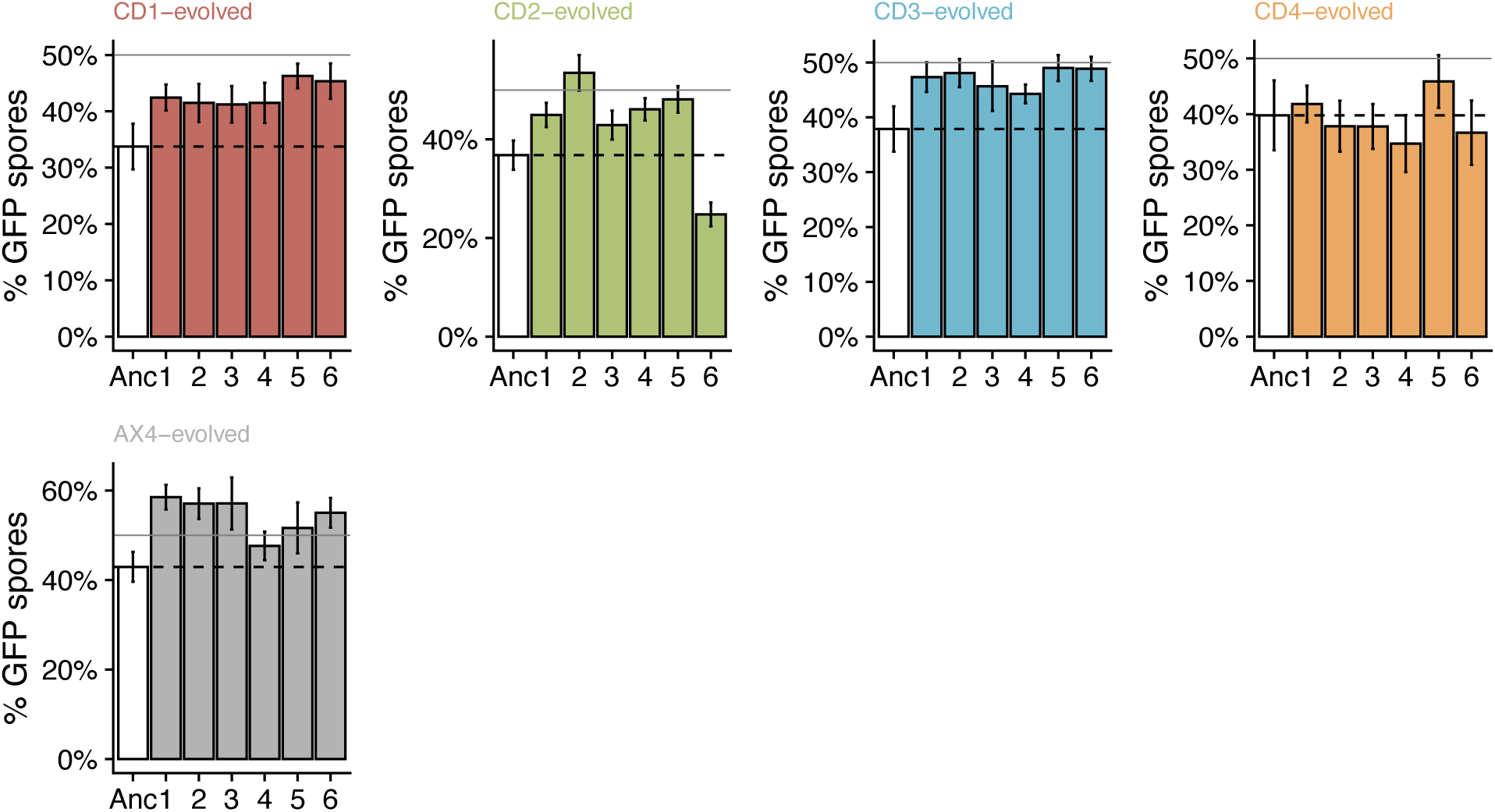
Cheater-resistance by single strains isolated from each evolved population. Each box shows the mean percentage of spores obtained by the indicated strain (ancestor or clones isolated from populations 1-6) following chimeric development with the cheater they evolved with. Dotted reference lines indicate the mean performance of the ancestor against that cheater. Solid lines indicate the expected percentage of GFP-labeled spores (=50%) if the tested strain were fully resistant to cheating. Except for CD4, most evolved isolates obtain a larger share of the spores than their ancestor did.

**Table 2.**
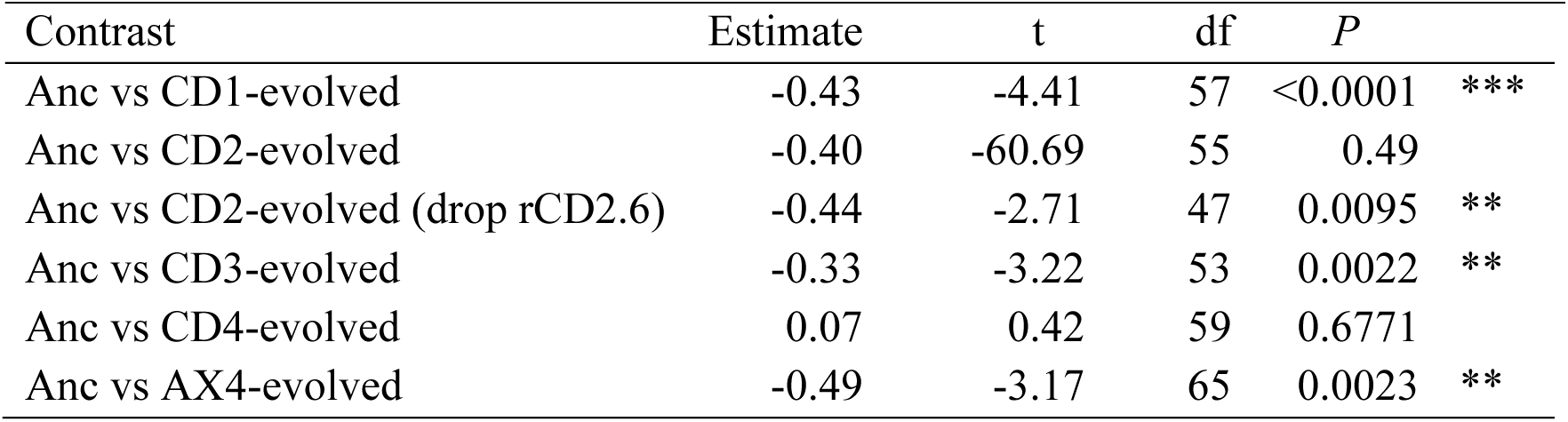
Contrasts comparing the performance of evolved isolates to their ancestor for each cheater (CD1-4) or the non-cheating control (AX4). Reported *P*-values are two-sided, but our hypothesis is one-sided (i.e., evolved populations should be better than the ancestor). CD2-evolved isolates are significantly better than their ancestor against the CD2 cheater once a single, poorly performing population is removed. However, the CD4-evolved isolates did not show significant evolutionary improvement.

### Do improvements against one cheater confer cross-resistance to other cheaters? Testing cheater specialization trade-offs

To test whether the improvements were specific to the targeted strain, we tested each evolved strain against all cheaters. Within this all-against-all design, there are two types of resistance: direct resistance, defined as improvements against the cheater that strain evolved with, and cross resistance, defined as improvements against cheaters that the strain has not experienced before (Fig. 1C). Cross-resistance thus arises as a correlated response to selection.

This design allows us to compare the magnitude of direct resistance to that of cross-resistance. We identified three different patterns we might see (Fig.5A): (1) complete cross-resistance, where strains do equally well against a given cheater, regardless of which cheater they experienced in the past; such a finding would implicate a general resistance mechanism and would be unlikely to lead to long-term coevolution, as resistance from prior cheaters confers strong resistance to novel cheaters. At the other end of the extreme are *direct costs of resistance*, where improvements against a focal cheater lead to losses of ability to resist other cheaters. Where direct costs of resistance arise, evolved strains should be better than their ancestor at resisting their focal cheater, but worse than their ancestor against novel cheaters. A third outcome is *opportunity costs of resistance*, where adapting to one cheater confers benefits against other cheaters, but these correlated responses to selection are not as large as the direct response. In this case, strains that adapt to one cheater are less well-adapted to some other (novel) cheater than strains that evolved with that cheater (CR<DR). Opportunity costs can lead to patterns of local adaptation, where the best-adapted strains are those that have encountered that cheater in the past, and, along with direct costs to resistance, can lead to continual coevolution because populations are never fully adapted to a novel cheater. Statistically, the key distinction is whether direct resistance is, on average, stronger than cross-resistance (DR>CR).

**Figure 5.**
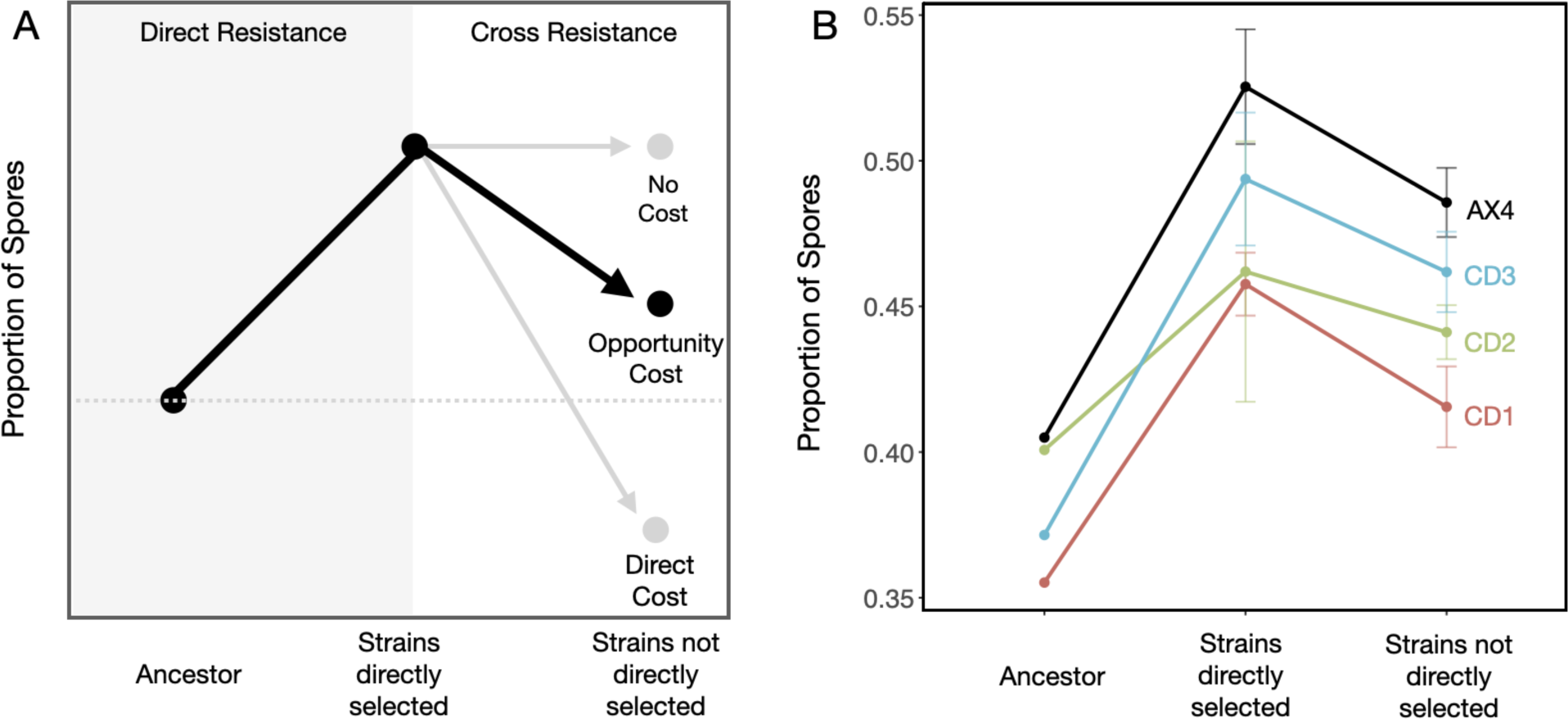
Local adaptation of cheating resistance. (A) Schematic of different hypothesized patterns. *No cost resistance (NC)*, where adaptations to one cheater are general and thus confer cross-resistance to unfamiliar cheaters. *Opportunity costs (OC)*, where strains that have adapted to one cheater (or partner) are less well-adapted to other cheaters (or partners) than strains that have had that opportunity. *Direct costs of resistance (DC)*, where evolutionary improvements against one cheater make those populations less well-adapted to another. Both OC and DC are examples of specialisation trade-offs. (B) Observed patterns of cross-resistance indicate opportunity costs. For CD1, CD2, CD3 and AX4, strains that have adapted with a particular social partner perform better with that partner than strains that did not evolve with that partner, indicating some cheater specialisation and that resistance is not absolute.

For populations evolving in response to three cheaters (CD1, CD2, and CD3) or the wildtype (AX4), we observe opportunity costs (Fig. 5B). In these treatments, strains that adapted to their partner strain also show improvements against other, unfamiliar strains — in other words, the evolved strains are also better than their ancestor against cheaters they were not exposed to. Thus, adapting to one cheater led to correlated improvements in performance against other cheaters.

However, although there was clear evidence of cross-resistance, cross-resistance was consistently weaker than direct resistance, indicated by the negative slope of the lines on the right side of Fig. 5B. This negative slope indicates that strains that evolve with a given cheater are, on average, better against that cheater than strains that evolved with some other cheater (or strain, in the case of AX4). This was true for CD1-, CD2-, CD3-, and AX4-evolved lines (emmeans contrasts, DR > CR; all *P*<0.05; Fig. 6). We exclude the CD4-evolved lines, as these strains did not show significant adaptation to their focal cheater. On average, ‘experienced’ strains—those exposed to a given cheater during their evolution—obtain 3.4% more spores than the ‘naïve’ strains do (those that experienced a different social partner during their evolution). Given that the average direct improvement was 10%, this result means that about 33% of the total improvement is strain specific. Thus, we see evidence of local cheater adaptation.

**Figure 6.**
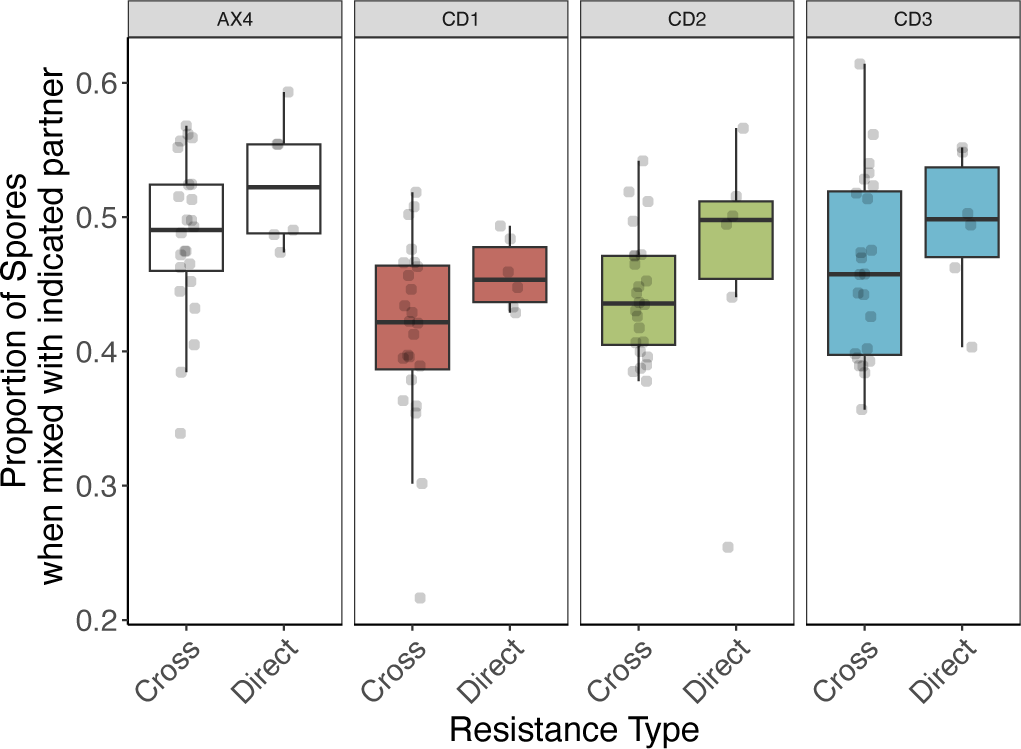
Cross resistance is weaker than direct resistance. Each plot shows the fraction of the spores that evolved strains obtain with the indicated ‘partner’ strain (either a CD cheater or AX4). “Direct” indicates the strains that were directly selected for performance with this partner, whereas “cross” indicates strains that were selected for performance with one of the other strains.

### Genomic Analyses of Cheater Resistance

To assess the genetic basis of cheater resistance, we used Illumina whole genome sequencing to identify mutations in a single isolate from each evolved line. We identified a total of 25 mutations in the 30 evolved lines (Table 1). The high AT content (77%) and low complexity of the *D. discoideum* genome make genome sequencing very challenging, for both traditional methods (see Eichinger et al. 2005) but for especially short-read sequencing methodologies such as Illumina. We therefore needed to adopt very stringent SNP filtering and manual review of each SNP, which means that we may miss some mutations. However, our stringent filtering criteria provide confidence that the mutations we have identified are correct.

Of the 25 mutations we observed, nine were missense, three were nonsense, six were synonymous, six were in intergenic regions (albeit close to the start of a gene in one case), and one was in an intron. Only 5 strains had more than one mutation; in all but one of these strains, we found only a single nonsense or missense mutation. Notably, we did not find any nonsynonymous substitutions in the rCD4 isolates, which is consistent with the lack of resistance in these strains. There were two possible examples of parallelism, where independent mutations are found in the same gene (Ostrowski et al. 2008). We found three different mutations in *mkcF*, a mitogen-activated protein kinase, each in a different line (rAX4.3, rAX4.5, and rCD2.3). Two of these mutations were nonsense mutations: one arose near the start of the gene, and the other towards the end, in the protein kinase domain. The third mutation was synonymous and also in the protein kinase domain. Little is known about *mkcF*, but an insertion mutant in this gene was enriched in two separate populations in a screen for ‘loser’ mutants — i.e., strains that become overrepresented in the anterior of the slug, a region containing the cells that have adopted the prestalk fate (Parkinson et al. 2011). In addition, the related gene, *mkcA*, is a negative regulator of sporulation (Shaulsky et al. 1996). Disruption of the *mkcA* gene also partially rescues the sporulation defects of *tagB*^-^ and *tagC*^-^ null mutants, suggesting a role in cell-fate decisions (Shaulsky et al. 1996).

In addition to parallel mutations in *mkcF*, we also observed two mutations in DDB_G0293998; however, both mutations arose in the same strain (rCD2.2), just three nucleotides apart and are likely to represent a single event. The rCD2.2 strain was the only isolate in which we identified more than one nonsynonymous or missense mutation (four total: 2 in DDB_G0293998, one in *tagA*, and one in *glcS*). This strain showed very strong resistance and is a possible cheater itself, obtaining >50% of the spores in chimeric co-development with CD2, the strain that exploited its ancestor (Fig. 4). Its mutations in *tagA* and *glcS* are also promising candidates for resistance alleles: the *tagA*-mutant has defects in cell-type specification and does not contribute to the prestalk-A region of the slug or to the terminal stalk when co-developed in the chimera with the wild-type strain (Cabral et al. 2006; Khare and Shaulsky 2010). The *glcS* null mutant has defects in stalk cell morphogenesis, resulting in small fruiting bodies, small spores, altered sensitivity to the stalk-inducing factor DIF, and reduced stalk cell vacuolization (Tresse et al. 2008). Finally, of the mutations in genes that have been subject to prior studies (e.g., *mkcF*, *nacA*, *med23*, *cdc5l, glcS*, *tagA*, *psiH*, and *grlA*), many are required for multicellular development and most have roles in prestalk differentiation or sporulation (Table S2; see also Glöckner et al. (2016)).

#### Life-history trade-offs

Local adaptation of evolving populations to strains they encountered recently suggests the possibility of lagging resistance, whereby resistance is always one step behind whatever new cheater mutation arises. It is thus one source of trade-offs, in that adaptation to any one cheater poses an opportunity cost for adaptation to a different cheater. A second type of trade-off is among different fitness components. We examined three fitness components that should all be under strong selection in this regime: doubling time during exponential growth, sporulation efficiency (the number of spores produced from a given number of starving cells), and spore germination efficiency (proportion of spores that produce amoebae). There were no significant differences among the selection regimes in doubling time (*χ^2^*=3.63, df=4, *P*=0.46; Fig. 7A) or sporulation efficiency (*χ^2^*=6.44, df=4, *P*=0.26; Fig. 7B). However, for germination efficiency, strains that evolved in the absence of a cheater (rAX4 strains) show better spore germination than the other evolved strains that co-developed with a cheater (*χ^2^*=15.5, df=4, *P*=0.004; Fig. 7C).

**Figure 7.**
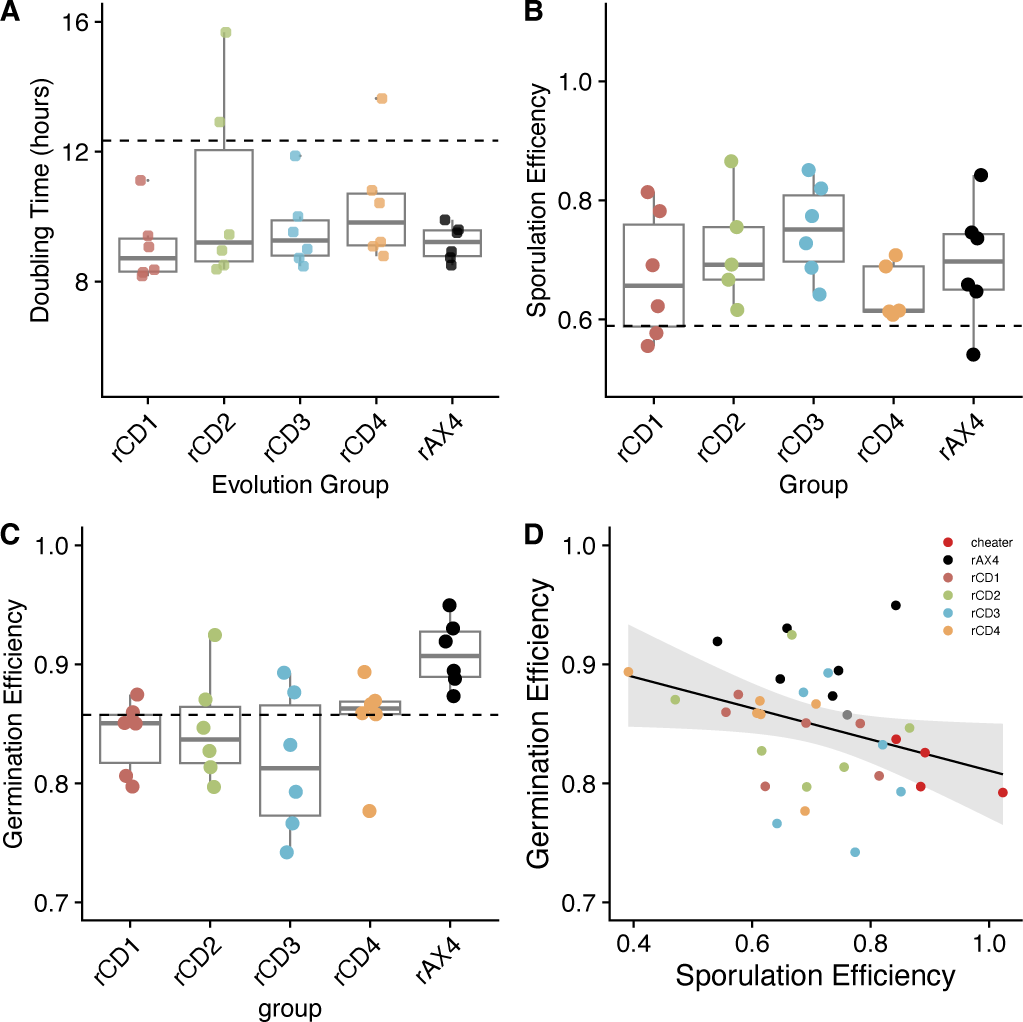
Does cheater resistance trade-off with improvements in other selected traits? **(A-C)** Each plot shows the performance of evolved isolates, relative to their ancestor for: (A) doubling time, (B) clonal sporulation efficiency, the fraction of starving cells that become spores, and (C) germination efficiency, the fraction of spores that germinate to produce amoeba. (D) Considering all strains together, there is a negative relationship between sporulation and germination efficiency (*r* = -0.35, df=33, *P*=0.04)

There are two possible explanations for this finding, both indicating a type of trade-off. One possibility is that same mutations that improve cheater resistance also make spore germination worse (antagonistic pleiotropy). An alternative possibility is that the mutations that confer resistance and germination improvements are different, but selection for resistance mutations impedes selection for germination improvements owing to clonal interference in those populations exposed to a cheater (clonal interference hypothesis). We return to the implications of these two possibilities in the Discussion.

To assess whether these two traits, cheater resistance and germination efficiency, trade-off we assessed their correlation across all of our strains. If pleiotropy can explain these results, then we expect to see a significant negative correlation. For cheater resistance, we used each strain’s grand mean spore fraction against all cheaters as a measure of its overall ability to resist cheating. There was no significant negative relationship between a strain’s dominance in the spores and its doubling time (*r* = 0.24, df = 28, *P* = 0.25) or germination efficiency (*r* = -0.22, *df* = 28, *P* = 0.25). However, resistance was strongly correlated with sporulation efficiency (*r* = 0.81, df=28, P<0.001), and across all strains (including the cheaters), sporulation efficiency is negatively associated with germination efficiency (*r* = -0.35, df = 33, *P*=0.04; Fig. 7D). Taken together, these results suggest that adapting to a cheater impeded adaptation to other aspects of the environment. Moreover, the negative relationship between resistance and sporulation and sporulation and germination suggests that spore number and spore quality may not be fully independent traits, supporting the hypothesis that the trade-off might be driven by antagonistic pleiotropy, whereby improvements in social traits come at a cost to other components of fitness.

**Table 1.** Genetic changes in evolved lines. Substitutions were identified using Illumina sequencing of one evolved clone from each experimental population. Details of SNP calling is provided in the methods section.

| Isolate | Chr | Position | Gene (or gene region) | Mutation | Mutation Type | AA Impact | Protein Length (aa) |
| --- | --- | --- | --- | --- | --- | --- | --- |
| rAX4.2 | 2 | 4522039 | DDB_G0274925-DDB_G0274459 | G ->A | intergenic |  | N/A |
| rAX4.2 | 4 | 3192636 | DDB_G0285411 | C ->A | nonsense | Ser1075* | 1438 |
| rAX4.3 | 1 | 4222660 | *mkcF* | C ->A | nonsense | Glu584* | 677 |
| rAX4.4 | 2 | 6906864 | DDB_G0276461-DDB_G0295773 | G ->T | intergenic |  | N/A |
| rAX4.4 | 3 | 4434038 | *nacA* | T ->A | synonymous | Ser101Ser | 160 |
| rAX4.5 | 1 | 4224088 | *mkcF* | G ->A | nonsense | Gln108* | 677 |
| rCD1.1 | 1 | 2942931 | *abnB-abnC* | A ->T | intergenic |  | N/A |
| rCD1.2 | 1 | 4871655 | *med23* | A ->G | missense | Lys1390Glu | 1662 |
| rCD1.4 | 3 | 1918447 | *cdc5l* | A ->T | missense | Leu792Ile | 800 |
| rCD2.2 | 1 | 547970 | *glcS* | C ->G | missense | Pro820Ala | 878 |
| rCD2.2 | 6 | 2502391 | *tagA* | G ->T | missense | Gly358Cys | 1752 |
| rCD2.2 | 6 | 3597053 | DDB_G0293998 | G ->C | missense | Arg79Thr | 152 |
| rCD2.2 | 6 | 3597056 | DDB_G0293998 | C ->A | missense | Thr80Asn | 152 |
| rCD2.3 | 1 | 4222592 | *mkcF* | A ->G | synonymous | Asn606Asn | 677 |
| rCD2.4 | 3 | 926995 | DDB_G0278471 | A ->G | synonymous | Thr489Thr | 527 |
| rCD2.4 | 5 | 2791834 | *psiH* | G ->A | missense | Val205Ile | 715 |
| rCD2.6 | 1 | 1537478 | DDB_G0268146 | T ->A | synonymous | Val253Val | 350 |
| rCD3.2 | 2 | 1560795 | DDB_G0272432 | G ->A | missense | Ser316Leu | 477 |
| rCD3.3 | 5 | 1354374 | DDB_G0288289 | A ->T | intron |  | N/A |
| rCD3.3 | 6 | 2613789 | DDB_G0293258 | G ->T | missense | Arg476Ile | 1192 |
| rCD3.4 | 5 | 4636778 | DDB_G0290815-DDB_G0290837 | G ->A | intergenic |  | N/A |
| rCD4.1 | 5 | 2177445 | DDB_G0288965-DDB_G0288967 | T ->G | intergenic |  | N/A |
| rCD4.1 | 5 | 2456867 | DDB_G0289195 | A ->G | synonymous | Glu35Glu | 296 |
| rCD4.2 | 2 | 782589 | *grlA*-DDB_G0271802 | G ->T | intergenic |  | N/A |
| rCD4.6 | 1 | 202668 | DDB_G0267458 | C ->T | synonymous | Leu484Leu | 678 |

## Discussion

In this study, we test the role of enforcement in maintaining cooperation in evolving populations of the social amoeba *D. discoideum*. Under repeated co-development with a cheater that contributes more than its fair share of spores, we observe significant improvements in the ability of the evolved populations to gain spores in co-development with the cheater. When evolved isolates were tested side-by-side for their ability to resist cheating by their focal cheater and cheaters that were unfamiliar to them, we found that there were correlated improvements in the ability to resist novel cheaters. However, this ‘off-target’ resistance is usually smaller in magnitude than direct resistance. Thus, we see specialization of resistors to cheaters – i.e., locally adapted genotypes.

Local adaptation can potentially promote an extended co-evolutionary arms race, where cheating selects for resistance, and imperfect resistance creates an opportunity for counter-evolutionary changes to emerge. This partial specialization to the local genotype could arise because the mutations themselves have these effects, or because mutations that are highly specific to one cheater arise in a genetic background with other mutations conferring general benefits to the environment. Thus, future work will be necessary to examine the different candidate mutations in defined genetic backgrounds.

We also tested whether life-history trade-offs might arise, such that resistors show enhanced resistance to their focal cheater, but this resistance comes at a cost with respect to evolutionary improvements in other selected traits. Indeed, we observe that strains evolved in the absence of a cheater genotype (rAX4) show enhancements in spore germination that were not seen in cheater-evolved lines.

Taken together, these findings demonstrate that cheating in social systems can be mitigated through counter-evolutionary changes to suppress it. We also found that improvements in the ability to prevent cheating evolve over short timescales and in response to genotypically distinct cheaters, indicating that there are plenty of evolutionary routes to suppression. However, at least over the timescale of this study, resistance was quantitative and almost never absolute. Imperfect resistance means that multiple resistance mutations may be required to fully counteract the advantage of cheating. In turn, the commensurately slow loss of cheating from populations will potentially allow greater opportunity for cheaters to evolve their own counter-changes to resist the resistance. Thus, the ready-yet-imperfect evolution of resistance is one mechanism that might help to maintain these different strategies and promote continued coevolution over time.

We also observed a second brake on resistance: it was at least partly specific to the cheater. In general, improvement against one cheater also enhanced resistance to other cheaters. Note that these general improvements may have been general to all cheaters or general to the evolution environment (including its abiotic features). Future work is necessary to sort out the specific genetic changes and their isolated impacts. However, we also observed a significantly higher ability to resist a cheater if the focal strain had been exposed to that cheater during its evolution. We refer to this phenomenon as an ‘opportunity cost’, in that adaptation to one cheater is a lost opportunity to adapt to another. In other words, resistors are locally adapted to the cheaters that have been encountered most recently. Diversity in cheating and resistance strategies might thus be maintained in geographically heterogeneous populations and may lead to the geographic mosaics of coevolution that are observed in other types of coevolving systems (Thompson 2005; Hollis 2012).

Finally, we observe one final cost of resistance: improvements against a cheater potentially impede adaptation to other aspects of the environment. In this case, populations evolved in the absence of a cheater show greater improvements in spore germination than populations evolved in their presence. There are two possible explanations for this finding. One is antagonistic pleiotropy—that the same mutations that enhance resistance also reduce germination. We see some weak evidence that this pattern could result from antagonistic pleiotropy, in that the strains with greater sporulation efficiency (which, in turn, was strongly correlated with resistance) showed worse germination. In other words, there may be a genetic trade-off between the number and quality of spores. This finding is supported by earlier studies with natural isolates of *Dictyostelium*, which showed that stronger dominance in the spores is associated with smaller spores, and smaller spore size itself was associated with reduced germination (Wolf et al. 2015). However, pleiotropy is not the only possible explanation for this finding. An alternative possibility is that the mutations conferring cheating resistance and enhanced spore germination are distinct. In clonal populations such as these, different mutations that are co-segregating in a population can interfere with each other’s fixation, a phenomenon known as ‘clonal interference’ (Hill and Robertson 1966; Gerrish and Lenski 1998). Indeed, if germination-enhancing mutations have smaller fitness effects than resistance-enhancing mutations, then the latter may fix at the expense of the former, at least initially. This, too, is a type of opportunity cost. In this case, the evolution of resistance slows adaptation to other components of the environment and creates a drag on the population’s ability to adapt to other features of the environment.

Taken together, our results demonstrate that counter-evolution of resistance is a plausible mechanism to counter cheating. It also addresses a long-term conundrum: if cheating can be so readily countered by resistance, why are cheating behaviors commonly observed? While one possibility is that these behaviors are simply transient in natural populations, genomic studies have suggested that the relevant genes may be experiencing balancing selection that maintains variation (Ostrowski et al. 2015). For the reasons we have shown here, resistance is likely to evolve, but it is also not a panacea. Imperfect resistance and costs of resistance can allow cheating to be sustained in populations, creating opportunities for prolonged coevolution. If so, then social conflict may itself be a potent driver of prolonged evolutionary change.

## Notes

### Competing Interest Statement

The authors have declared no competing interest.

