## Supplement for "Costs of resistance limit the effectiveness of cooperation enforcement"

1. Supplemental Methods
2. Additional CD4 Analyses
3. Figures S1 and S2
4. Tables S1 and S2
5. Literature Cited

**Supplemental Methods**

*Whole-Genome Resequencing and Variant Calling* - Genomic DNA was prepared using phenol-chloroform extraction from axenic cell cultures. To obtain nuclei, cells were washed and resuspended in 15 ml nuclei buffer (40 mM Tris-HCl, pH 7.8, 6 mM $MgCl_2$, 40 mM KCl, 0.1 mM EDTA, 5 mM DTT, 1.5% sucrose, 0.4% IGEPAL) and incubated on ice for 10 minutes with several rounds of pipetting, followed by centrifugation at max speed at 4C to pellet the nuclei. Following removal of the supernatant, the nuclei pellets were stored at -80 °C until DNA extraction. To purify DNA, nuclei were thawed on ice and resuspended in 100 µl of 0.1 M EDTA. The following were added sequentially while incubating at 60 °C: 450 µl STE solution (10 mM Tris-HCl, pH 8, 10 mM EDTA, 400 mM NaCl), 50 µl 10% SDS, 10 µl of 10 mg/µl Proteinase K. The solution was incubated for 1 hour at 60 °C. Following incubation, the solution was mixed with 500 µl of phenol:choloroform:isoamyl alcohol by inverting and spun at top speed for 10 minutes. The aqueous phase was transferred to a new tube and the process was repeated until no interface was observed. The supernatant was then treated with one round of chloroform extraction, followed by ethanol precipitation (2X volume of 100% EtOH and 1/10 volume of 3M NaOAc). The pellet was air-dried and then resuspended in Tris-EDTA (pH=8) with 100 µg/ml ribonuclease A (RNase A) and incubated overnight at 4 °C. Following RNase treatment, phenol:chloroform extraction and ethanol precipitation was repeated to remove the RNase.

Libraries were prepared using the Illumina NexteraXT kit according to the manufacturer's protocol, except that reaction volumes were reduced to 0.25× before size selection using

SPRIselect beads (Becker Coulter B23319) for a target size of 300-500 bp. Raw sequencing reads were mapped to the chromosomal sequences of AX4 (GFF files dated 30 November, 2016, available at: http://dictybase.org/download/gff3/) using BWA-MEM (Li and Durbin 2009). The AX4 genome contains a large (~750 Kb) duplication within chromosome 2, which was masked from the reference for the alignment. Duplicate reads were marked using Picard 2.18.13 MarkDuplicates (http://broadinstitute.github.io/picard). Variant calling, genotyping, and hard filtering was performed with GATK4.0.11.0 (McKenna et al. 2010) following GATK Best Practices (Depristo et al. 2011; Van der Auwera et al. 2013). Base quality score recalibration was performed on a joint-call cohort and final variant calling was applied to individual samples. Variants were called using GATK4 Haplotype Caller in GVCF mode. After initial variant calling, filtering thresholds were applied to remove variants likely to be false positives. These thresholds were set for eight statistics which describe each variant and provide evidence on the likelihood of each being a true variant. The threshold for Read Position Rank Sum (ReadPosRandSum) was determined using Tukey's method to identify outliers scored above and below 1.5×IQR (inter quartile range). QualitybyDepth (QD), Mapping Quality (MQ), StandOddsRatio (SOR), MappingQualityRankSum (MQRankSum) were set according to the GATK4 hard filtering recommendations (available at: https://software.broadinstitute.org/gatk/documentation/article.php?id=6925).

Variants were filtered by the following criteria: (QD > 2.0, FS < 60, MQ > 40, SOR < 7.0, MQRankSum > -12.5, ReadPosRankSum > -8). A maximum depth filter of 2x mean per sample was applied to reduce spurious calls, while the minimum depth required was three. Additionally, variants with a depth at or above mean + 3X sqrt mean and Qual < 2X mean depth were removed (see Li 2014). To further reduce the likelihood of false positives, we required that at least 80% of the reads covering a site showed the alternate allele. We also required that at least one strain in the cohort showed strong evidence for the reference allele (>80% of reads covering the site show the reference allele). Annotations were added to variants using SnpEFF v4.0 (Cingolani et al. 2012)with AX4 GFF3 file generated November 30$^{th}$, 2016 (available at: http://dictybase.org/download/gff3/).

**Additional CD4 analyses.** Although most populations and evolved isolates showed improved ability to obtain spores when co-developed with their targeted strain, the isolates from the rCD4 populations showed a different pattern. Analyses of the evolved populations showed improvements against CD4 (Fig. 3), but when we isolated single clones and tested their ability to form spores, they did not show significant improvement, and the mean fraction of spores they obtained when co-developed with CD4 was less than that of the ancestor in all cases (see Fig. 4). On one hand, this result is not too surprising: evolving populations can be genetically diverse, and any given clonal isolate will not necessarily be representative of the population as a whole. Clonal isolation also involves a single-cell bottleneck, which can lead to the acquisition of deleterious mutations. Finally, if there are genotype-by-genotype (G x G) interactions within a population, then a clonal isolate may have a different fitness owing to the different genetic context in which it is being tested.

While these explanations can explain low fitness in any given isolate, it was nonetheless surprising to see that all six isolates performed worse than their ancestor against CD4, the strain they evolved with. Further investigation pointed out several mitigating factors. First, the lack of resistance observed for the rCD4 isolates appear to be caused more by *higher* fitness of the ancestor (to which they are being compared) than *lower* fitness of these isolates (i.e., compare the ancestral values from Fig. 3 and Fig. 4). When we looked at why the ancestor had higher fitness (when co-developed with CD4) in these assays, we found that the starting frequencies of the cells deviated substantially from 50-50, with the ancestor starting in some cases at a frequency >0.7. This meant, in turn, that the ancestor came out at a higher frequency in the spores, too. (Note that, on average, two competitors start at 50-50, but the exact starting frequency can vary across replicates owing to counting and experimenter error.)

To ensure that deviations in starting ratios were not impacting our results, we added a filter that retained data points only if the starting percentages of cells were between 0.4 and 0.6. With this filter in place, the ancestor vs. CD4 estimate was now similar to our previous estimates (see Figure S2; compare to Fig. 2 and Fig. 3) and only two populations still show a mean performance lower than that of the ancestor. Nevertheless, as a group, the rCD4 isolates were still not significantly better than their ancestor against CD4 (emmeans contrast, rCD4 vs. ancestor = 0.03, $P$=0.90). Thus, the isolates are not significantly resistant. However, these analyses also indicate that we do not see any strong evidence that the rCD4 isolates picked up deleterious mutations during the cloning process. Further supporting these conclusions (lack of resistance, as well as lack of deleterious mutations), we also failed to identify any nonsynonymous mutations in these isolates.

**Figure S1. Publicly available information on the timing and cell-type expression patterns of gene disrupted in CD strains. (A)** Expression time course of the affected genes during multicellular development in the standard lab strain, AX4. The x-axis shows the number of hours since starvation commenced, and development is completed at 24 hr. The y-axis shows the normalized number of reads mapping to that gene. (B) Degree of cell-type expression bias for genes disrupted in CD strains. The x-axis is the log2 fold coverage ratio of prespore to prestalk cells. The y-axis is the -log10 of the false discovery rate; large values (>2) indicate significantly biased expression differences between these two cell types. Data and figures produced by DictyExpress[1] and are based on data collected by Rosengarten et al.[2] (A) and Parikh et al.[3] (B).

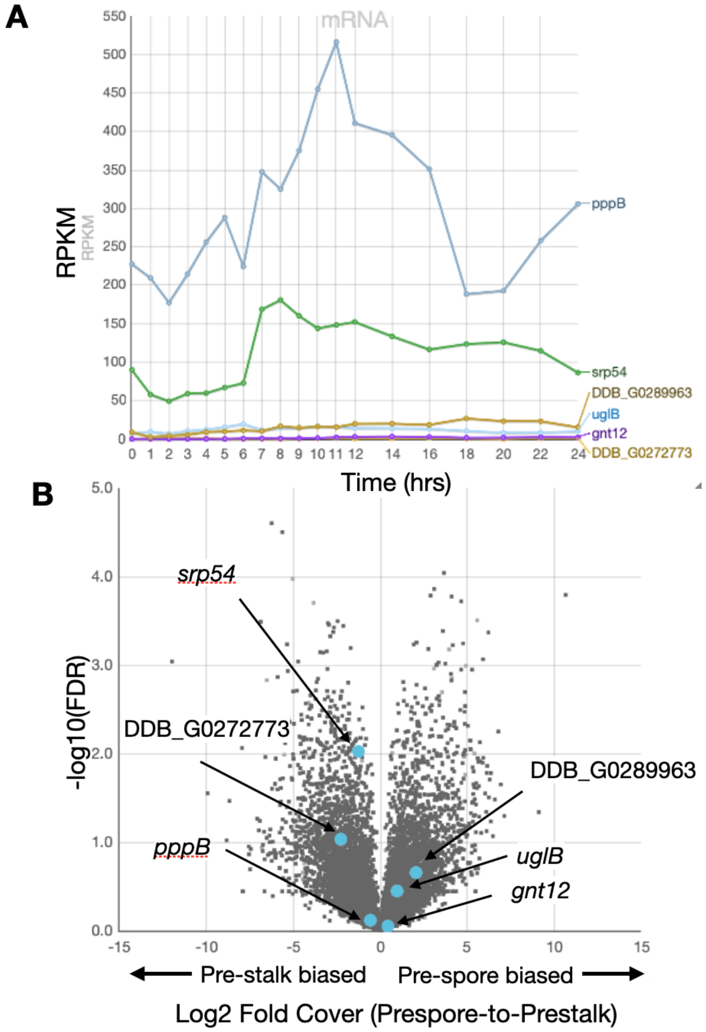

**Figure S2. Resistance of clonal isolates from CD4-evolved populations.** Each box shows the mean percentage of spores obtained by the indicated strain (ancestor or clones isolated from populations 1-6) following chimeric development with the cheater they evolved with. Dotted reference lines indicate the mean performance of the ancestor against that cheater. Solid lines indicate the expected percentage of GFP-labeled spores (=50%) if the tested strain were fully resistant to cheating. In contrast to our earlier analysis (Fig. 4), in this analysis, we retained only data points where the starting percentage of the two strains was between 0.4 and 0.6. However, the rCD4 isolates are still not significantly better than the ancestor at obtaining spores when co-developed with CD4.

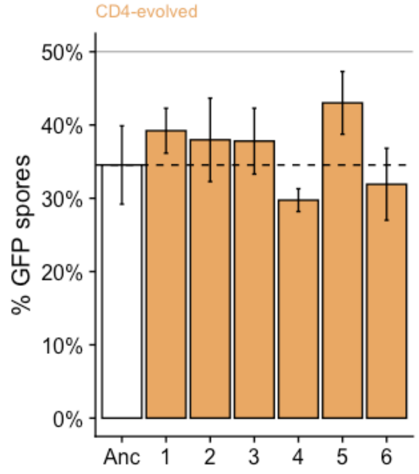

**Table S1. Locations of integration of the REMI plasmid in the cheater strains, CD1-CD4.** These sites were identified by plasmid rescue (by Chris Dinh, BCM) and/or whole genome Illumina sequencing (by Michael Miller). Annotations retrieved from dictybase.org (18 May 2023), and the link to the Dictybase webpage is provided.

<table>
<tr><th>Strain</th><th>Chromosome: position</th><th>Position in gene</th><th>Gene Product / Description</th></tr>
<tr><td>CD1</td><td>2:2160158</td><td>intergenic</td><td>DDB_G0272773 -> | <- *uglB*<br>*uglB* = uracil glycosylase</td></tr>
<tr><td>CD2</td><td>2:6109796</td><td>intergenic</td><td>*pppB* <- | <- *srp54*.<br>*pppB* = protein phosphatase P family protein; *srp54* = Component of the signal recognition particle (SRP), involved in targeting of nascent secretory proteins to the ER.</td></tr>
<tr><td>CD3</td><td>1:1578392</td><td>489 (exon 2)</td><td>*gnt12*, putative glucosyltransferase</td></tr>
<tr><td>CD4</td><td>5:3511091</td><td>DDB_G0289963</td><td>No annotation</td></tr>
</table>

**Table S2. Information about gene products, related genes, null phenotypes, and timing of expression for genes that were mutated in evolved lines.** The genes below are a subset of those listed in Table 1 for which we could find published data about their function on Dictybase[4]. Timing of expression is based on data from Parikh et al. 2010[3]. None of the genes showed significant prespore-to-prestalk bias in expression (based on analyses of data from Parikh et al. 2010[3] using DictyExpress[1]; data not shown).

| Gene Name | Gene Description (from dictybase[4]) | Null phenotype (from dictybase[4]) | Timing of expression (vegetative, developmental, both, or not expressed[3]) |
|---|---|---|---|
| *cdc5l* | Cell division cycle 5-like; contains two Myb DNA-binding domains; ortholog of human CDC5L and yeast CEF1 | Not available | Both |
| *glcS* | Glycogen synthase | Aberrant stalk morphogenesis, reduced vacuolization, decreased fruiting body size, decreased glycogen level, decreased spore size, aberrant cell morphology[5] | Both |
| *med23* | Putative mediator complex subunit 23 | Aberrant culmination, aberrant oscillatory cAMP signaling | Both |
| *mkcF* | Similar to *Dictyostelium mkcA, mkcB,* and other mitogen-activated protein kinases (Ste20/PAK family); *mkcA* is a negative regulator of sporulation[6] | Not available. However, *mkcF* insertion mutants were isolated in a screen for 'losers'—mutants that show prestalk preference[7]. Similar to genes *mkcA* and *mkcB:* the *mkcA* null mutant shows decreased prestalk expression and decreased sporulation relative to the wild-type. It is a partial suppressor of the *tagB*$^-$ sporulation defect[6]. An *mkcB* insertion mutant showed decreased fruiting body size[8]. | Both |
| *nacA* | Putative nascent polypeptide-associated complex alpha subunit | Not available | Both |
| *psiH* | Similar to prespore inducing factor PsiA, a secreted protein that regulates both prespore and prestalk cell differentiation | Not available; *psiA* null mutant shows abolished prespore cell division, precocious prestalk gene expression, abolished prespore gene expression[9,10] | Both |
| *tagA* | ABC transporter B family protein, serine protease | Decreased spore cell differentiation, decreased spore germination, decreased sporulation, delayed culmination, aberrant fruiting body morphogenesis, increased number of mound tips, increased prestalk A differentiation, abolished cell migration to prestalk region | Both |
| DDB_G0285411 | Ankyrin repeat-containing protein cyclin-like F-box containing protein | | Both |
| DDB_G0293998 | No annotation | | Developmental |
| DDB_G0272432 | No annotation | | Developmental |
| DDB_G0293258 | Putative myotubularin phosphatase Mtm3 | | Both |
